# Primate visual cortex spontaneously computes the beauty of objects

**DOI:** 10.64898/2026.07.07.736498

**Authors:** Philipp A Schumann, Rico Stecher, Martin N Hebart, Daniel Kaiser

## Abstract

During everyday vision, beauty is a spontaneous experience. Perceived beauty varies substantially across object categories: We find puppies more beautiful than moths and roses more beautiful than potatoes. Here, we provide evidence that early processing in human and non-human primate visual cortex already reflects perceived beauty, even without deliberate judgments of beauty. First, in a combined EEG-fMRI analysis, we show that beauty ratings for hundreds of object categories from the THINGS database are predicted from neural responses measured in separate groups of human participants viewing the same images without explicit beauty judgments. Beauty-related representations emerged in occipital visual cortex within 100ms and peaked around 130ms. Second, multi-unit activity in macaque visual cortex also predicted human beauty judgments, despite the animals’ highly limited knowledge of the objects. These results indicate a perceptual basis for the beauty of visual categories, which may underlie spontaneous preference formation in everyday situations.

**Teaser:** Early visual areas in both human and non-human primates rapidly compute the beauty of real-world objects

## Introduction

The experience of beauty is an integral part of our everyday lives (*1, 2*). For example, when walking down a street, we may admire a beautiful building or a stunning cloud formation (*3*). Within coarse-scale visual domains, such as architecture, faces, or artworks, there are examples that are consistently perceived as more or less beautiful. However, everyday visual objects themselves also differ vastly in their aesthetic appeal: We find roses more beautiful than raisins and flamingos more beautiful than vultures. Here, we investigate how the beauty of objects is fundamentally determined by perceptual computations in the visual cortex.

Some previous research suggests that aesthetic experience requires thought (*4*) and therefore depends on intentional higher-level cognition linked to neural responses in orbitofrontal areas (*5*) and the Default Mode Network (*6*). Contrasting this view, numerous EEG studies have reported rapid beauty-related responses, sometimes starting as early as 100ms after stimulus onset, indicating that representations of beauty first emerge in visual cortex (*7-10*). More recent computational approaches demonstrate that even artificial neural networks with orthogonal training objectives like categorization are capable of predicting human beauty judgments (*11-15*), suggesting that visual beauty is firmly grounded in the analysis of visual features.

Yet, previous investigations only assessed the neural correlates of perceived beauty across exemplars of one or few categories, often failed to account for alternative explanations related to related concepts like emotional valence, and provided a characterization limited to the spatial or temporal domain. Furthermore, most studies employed explicit beauty-rating tasks while measuring beauty-related neural responses, leaving it unclear how beauty arises spontaneously from cortical activations that emerge incidentally during visual processing. Here, we provide a strong test of such a representation by (1) delineating uniquely beauty-related representations across a very broad range of visual categories, and (2) by assessing how the beauty of visual categories can be predicted from visual responses that occur incidentally, without a beauty-related task. If such incidental visual responses indeed predict beauty on the category level, we predict that they do so during early processing in the visual cortex. In addition, we hypothesize they are fundamental enough to not only occur in humans, but even in a non-human primate model.

In our study, we predicted human beauty ratings for hundreds of object categories from brain responses in a set of participants that merely viewed these images in a task regime unrelated to beauty. We used EEG, MEG and fMRI responses in humans (*16, 17*) and performed multimodal fusion analysis (*18, 19*) to resolve the neural representations of beauty in both space and time. To validate and extend our results across species, we further predicted beauty ratings from visual cortex activity in non-human primates that viewed the same images (*20*).

Our results show that early responses within 100ms of processing, localized to the visual cortex, predict the beauty of visual categories, demonstrating a spontaneous, rapid representation of beauty that emerges in parallel to object recognition and in the absence of a beauty-related task. Moreover, we show that neural responses to the same objects in the macaque visual cortex also predict human beauty ratings, revealing that early visual responses predict beauty even for non-human primates. Together, these findings establish a perceptual neural basis for the perceived beauty of natural object categories, which may form the basis for the spontaneous appreciation of beauty in everyday situations.

## Results

To reveal the neural correlates of beauty across many visual categories, we employed an extensive sampling approach, based on the THINGS database (*21*), which contains photographs of 1,854 object categories. In an online experiment, observers (N=3,750) rated the beauty of 7 different images for each of the 1,854 categories, with each image rated by 24 to 36 raters (M=30.51). Collapsing across the individual images, we obtained beauty ratings for each category, which were subsequently related to neural responses to the object categories, recorded in experiments unrelated to beauty.

### Rapid responses in human visual cortex predict the beauty of visual categories

We related beauty ratings to spatially and temporally resolved human neuroimaging data, using human fMRI (*17*) and EEG (*16*) recordings (see Fig. 1 A), acquired for a subset of 634 out of the 1,854 object categories. During the EEG and fMRI sessions, participants viewed 12 example images for each category while performing a simple oddball detection task that was unrelated to beauty (see Methods). Notably, the EEG data was recorded under a rapid presentation regime, which likely prevented participants from making slow and deliberate judgments of beauty. This experimental regime allowed us to assess whether neural representations that arise spontaneously and are unrelated to beauty judgments can still predict the beauty of object categories as assessed by our online observers.

**Figure 1.**
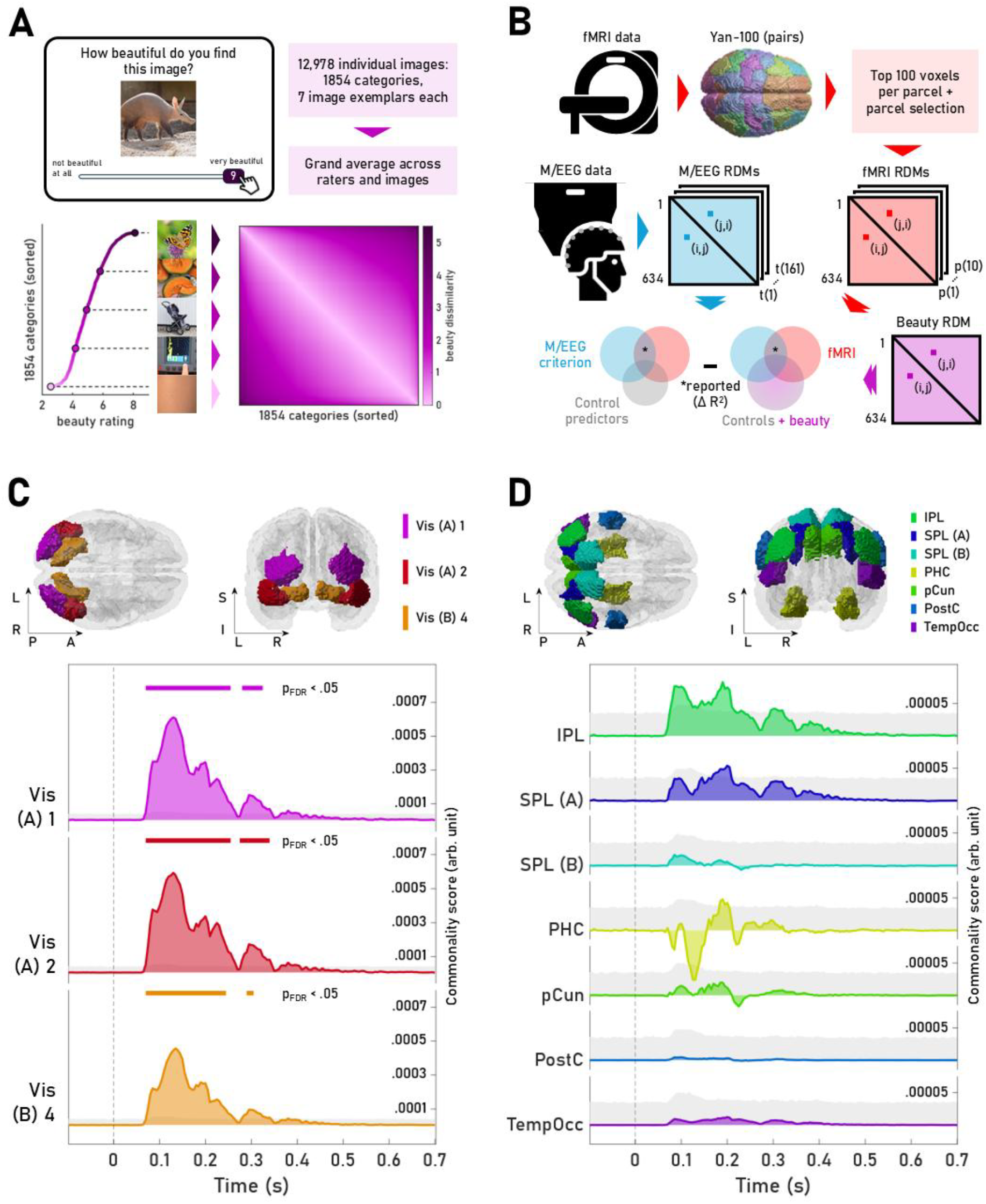
EEG-fMRI fusion analysis reveals a spatiotemporal representation of beauty. A) Human ratings of category-level beauty were collected in an online task and averaged across observers and exemplar images. The most beautiful category was butterflies and the least beautiful category was skin (see Supplementary Fig. 3 for the Top 10 most and least beautiful categories). Pairwise absolute differences in beauty ratings were then arranged into a Representational Dissimilarity Matrix (RDM). B) We conducted an RDM-based commonality analysis by integrating (1) fMRI data (parcellated and preselected), (2) EEG data (across time), and (3) beauty ratings (while controlling for other object property ratings), across the shared subset of 634 object categories (200,661 pairwise comparisons). The Venn diagrams illustrate two semipartial correlations (The variance partitioning schema is for illustration purposes only. See Methods for details). We report the difference between two squared semipartial correlation coefficients as the commonality score. C) Occipital visual parcels (visualized in MNI space) and stacked ridge plots displaying the time-resolved commonality scores per parcel. Significant commonalities (FDR-corrected p<.05, permutation tests, indicated by horizontal bars) emerged well before 100ms after stimulus, with a peak at 130ms, and sustained significance beyond 200ms. D) In areas outside the visual cortex, commonalities were not significantly greater than expected when correcting for multiple comparisons across time (light grey shaded area: max permutation statistic capped at its 95^th^ percentile). Abbreviations: Vis: Visual, IPL: Inferior Parietal Lobule, SPL: Superior Parietal Lobule, PHC: Parahippocampal Cortex, pCun: Precuneus, PostC: Postcentral, TempOcc: Temporal Occipital.

To obtain a spatiotemporal mapping of beauty-related representations, we performed model-based EEG-fMRI fusion analysis (*19, 22, 23*). Analogous to representational similarity analysis (RSA), we first computed representational dissimilarity matrices (RDMs) capturing pairwise dissimilarities of EEG responses to the stimuli at every time point and then averaged them across participants. These RDMs provided a temporally resolved view across the visual processing hierarchy. Second, we computed pairwise dissimilarities in fMRI responses in each of 50 homotopic bilateral brain parcels (*24*), which we again averaged across participants. The resulting RDMs provided a spatially resolved view across the whole brain. Bilateral parcels were chosen as we did not expect strong lateralization effects, given the stimulus diversity. Third, we computed the pairwise dissimilarities in beauty ratings averaged across all exemplars for each category. The resulting RDM provided a measure of how similar each category was to every other category in its perceived beauty. All RDMs had 200,661 unique entries, i.e., pairwise comparisons of 634 categories. We utilized commonality analysis (*25*) to calculate the tripartite shared variance between EEG, fMRI and beauty ratings, extracting the timepoints and parcels where a portion of the representational similarity between EEG and fMRI is driven by beauty.

Critically, to ensure that the resulting commonality coefficients are uniquely attributable to dissimilarities in beauty, rather than other judgments correlated with beauty, all models accounted for dissimilarities in a set of 12 other property ratings for all categories: Manmadeness, naturalness, preciousness, pleasantness, cuteness, arousal, animacy, heaviness, as well as the abilities to move, be moved, grasped, and held. These ratings were taken from the THINGSplus dataset (*26*). For each of these properties, we generated an RDM based on pairwise rating differences, which we partialled out in all analyses.

We report commonality score time courses, epoched between -100ms and 700ms relative to stimulus onset, for a set of 10 bilateral parcel pairs from the Yan atlas (*24*). These parcels were selected a priori, based on an initial RSA solely on the fMRI data, in which they yielded significant correlations between regional fMRI RDMs and beauty RDMs. Parcels included regions in the occipital visual cortex, the dorsal attention network (*27*) as well as the default mode network (see Fig. 1 C & D, for a detailed report see Supplementary Table 1). Significance of the commonality coefficients was assessed using a permutation test at each time point and False Discovery Rate (FDR) corrections (*28*) across time. For this purpose, we developed a novel permutation testing routine that isolates the effects of the critical predictor beauty while accounting for spurious effects that are caused only by the collinearity of the regressed variables (see Materials and Methods). This provides a methodological advance over existing similarity-based fusion approaches for more appropriate inferences on variance partitions.

We found a significant and relatively sustained commonality effect of beauty in all three visual parcels (Fig. 1 C). Remarkably, the onset of these representations fell within the first 100ms of processing across all parcels, indicating that rapid neural responses, even during a beauty-unrelated task, correlate with beauty judgments. These parcels, located in the occipital visual cortex, show a comparably prominent and temporally similar commonality time course, which is characterized by a sharp onset at 65ms, as well as a strong early peak at around 130ms after onset (see Fig. 1 C). After this early peak, the visual cortical areas show a decreasing commonality score, albeit continued significance up to 250ms post onset, and further significant effects around 300ms. It is worth highlighting that these robust, relatively prolonged beauty-related activations occurred despite the rapid presentation regime in the EEG experiment (200ms stimulus onset asynchrony). Commonality scores in all other parcels (see Fig. 1 D) were an order of magnitude lower than in occipital visual parcels. They also displayed a less homogeneous pattern: Areas located in the parietal cortex, such as portions of the inferior and superior parietal lobule, showed an early onset and a global peak around 200ms. Yet, none of these commonality coefficients reached statistical significance after multiple-comparison correction.

We performed two supplementary analyses solidifying our results. First, we investigated whether our results truly reflect beauty on the category level, rather than the beauty assigned to individual images. To this end, we re-ran the EEG-fMRI fusion analysis using only the exemplar images shown in the EEG and fMRI study that were not rated for beauty in the online experiment. The results in occipital visual parcels remained significant (see Supplementary Figure 1 A), indicating that the representations are indeed driven by the beauty of object categories rather than individual exemplar images. Second, we tested the reproducibility of our results across neuroimaging methods. To this end, we additionally performed an MEG-fMRI fusion analysis using MEG data recorded for the same object images (*17*). This analysis yielded a notably similar pattern, showing an early and reliable neural representation of beauty (see Supplementary Figure 1 B).

Together, these results reveal a rapid representation of the beauty of object categories in the visual cortex, spatially and temporally co-occurring with cortical responses related to object recognition (*16, 17, 22, 29*). Representations of beauty emerged within the first 100ms of processing and were sustained beyond 200ms. The significant and much larger commonality coefficients in occipital visual areas, in contrast to the parcels outside the visual system, suggest that visual object processing is immediately reflective of beauty, in particular early processing in the visual cortex.

### Responses in monkey visual cortex predict human beauty judgments

Our human neuroimaging results suggest that category-level beauty is fundamentally reflected in visual cortex responses that emerge spontaneously during object perception. Humans, however, learn to associate certain object categories with their beauty, and visual representations may be shaped by those associations. For instance, we may repeatedly see rainbows, have a pleasurable feeling when seeing them, and talk about their beauty. The association with beauty may shape their representation in visual cortex. Alternatively, there may be something fundamentally sensory about the beauty of rainbows, with their visual features acting as a true precursory of beauty, even in the absence of such associations. If there existed such a visual precursor for perceiving beauty, we should be able to predict human beauty judgments from cortical responses in non-human primates as well, given their highly similar visual object processing architecture (*30, 31*). A successful prediction from monkey visual cortex would further corroborate that the effects are largely independent from human associations with an object’s beauty: After all, monkeys will not have encountered, and appreciated, most of the object categories in the THINGS repertoire.

Here, we analyzed multi-unit activity (MUA) recorded from regions V1, V4, and IT of two macaque monkeys who viewed 12 exemplars of all 1,854 object categories (*20*) (see Fig. 2 A). Specifically, we performed RSA on normalized MUA around the peak in the MUA time courses (75ms in V1, 100ms in V4, and 125ms in IT) (*20*). We created a MUA RDM for each of the three regions by correlating the normalized MUA across recording sites in each region between every possible pair of stimuli and subtracting the resulting correlations from 1, yielding 1,854 by 1,854 RDMs (∼1.72M pairwise comparisons) for each region and both animals. We then correlated these RDMs with the beauty RDM constructed from pairwise dissimilarities in human beauty ratings while, similarly to the EEG-fMRI fusion, all while partialling out the 12 other property rating RDMs.

**Figure 2.**
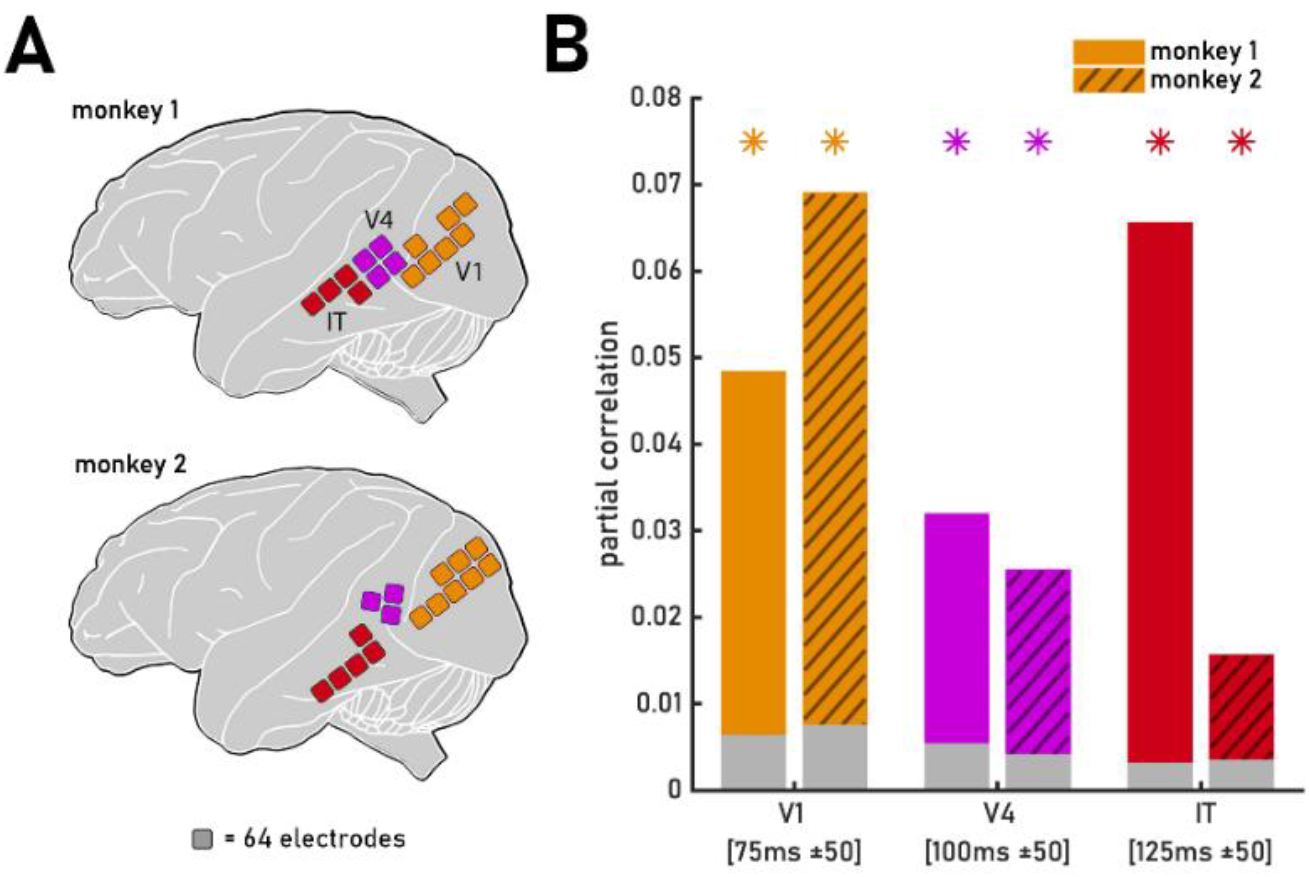
Multi-unit activity (MUA) in monkey visual cortex predicts human beauty ratings. A) We analyzed MUA from two macaque monkeys with electrodes in V1, V4, and IT. The animals viewed 12 exemplars of all 1,854 object categories in the things database. Illustration adapted from (20). B) Representational similarity analysis on peak MUA patterns across electrodes revealed that pairwise dissimilarities in MUA patterns and beauty ratings were significantly correlated with dissimilarities in beauty ratings (with control ratings partialled out). This relationship emerged across regions and across both animals, indicating robust prediction of human beauty ratings from monkey visual cortex. Asterisks indicate p<0.05. Shaded bars represent 95% confidence margins from a permutation test.

For both animals, we found significant correlations (permutation tests, FDR-corrected for multiple comparisons) between the beauty RDM and the MUA RDMs in V1, V4, and IT (see Fig. 2B). Early neural activity in monkey visual cortex (extracted within the first 150ms of processing in all three regions) thus predicts human beauty judgments, suggesting that rapid and mandatory computations in visual cortex capture perceived beauty. The rapid onset of neural activations related to perceived beauty is also apparent in analyses of time-resolved MUA (see Supplementary Fig. 3). Note that differences across regions and animals may relate to differing numbers of electrode contacts across regions and different electrode placement across animals. In sum, these results support the conclusion that visual representations in primate cortex spontaneously reflect the beauty of object categories, suggesting a shared ancestry for the fundamentally visual representations that predispose the human experience of beauty.

## Discussion

We show that the beauty of object categories is coded in early neural responses in the visual system. By modelling neural representations in temporally and spatially resolved brain responses, validated across species, our results convergently suggest a perceptual basis for the beauty of visual objects, even in the absence of a beauty-related task regime. This is consistent with the remarkable predictive power of computer vision models for beauty judgments, even in the absence of beauty-related training regimes (*11-14, 32*). Such representations of beauty may function as a perceptual basis for aesthetic experiences with visual stimuli, enabling the spontaneous, sometimes surprising, experiences of beauty in everyday life.

The neural representation of visual beauty during perceptual processing aligns well with theories in visual aesthetics (*5, 32-39*), which posit that the analysis of perceptual attributes and its corresponding activity in sensory cortical areas is critical for aesthetic appreciation. It is further consistent with previous findings targeting the neural correlates of beauty within stimulus categories like faces and scenes, with early EEG responses predicting within-category beauty (*8-10, 40-43*) and visual features extracted by DNNs predicting human beauty ratings (*11-14, 44*). While many theories in visual aesthetics stress the importance of cognitive factors and meaningful engagement with a stimulus (*4, 6, 32, 45, 46*), our results complement these conceptions by showing that early visual responses may serve as a precursor to the aesthetic evaluation of object categories, even in the absence of explicit beauty judgments. These representations of category-level beauty in the visual cortex thus emerge somewhat independently of the task, in line with previous findings of a task-independent evaluation of symmetry in abstract patterns (*43, 47*) and consistent with recent EEG work showing no substantial difference in the emergence of representations related to scene beauty between an explicit beauty judgment task and an unrelated judgment (*48*). Together, obtaining our results in absence of a beauty-related task and under a rapid presentation regime in the EEG experiment suggests that the neural correlates of beauty partly originate from incidental processing in visual cortex that occurs without deliberate evaluation.

While occipital, occipito-temporal and occipito-parietal regions have been implied in the representation of visual beauty by several studies (*49-53*), others highlight the importance of regions outside the visual system, including the default mode network (DMN) (*6, 54, 55*), frontal brain areas (*6, 13, 56, 57*). In our EEG-fMRI fusion, we did not find significant effects in such non-visual regions. This is consistent with the view that regions like the DMN are linked to the deliberate evaluation of a stimulus, for instance to assess its relevance in relation to the self (*55*). In our study, with a beauty-unrelated task and a rapid presentation regime, only the spontaneous perceptual foundations, and potential triggers, of such later and deliberate evaluations may emerge.

The spontaneous representation of beauty during visual perception is reminiscent of correlates and representations of emotion in the visual cortex (*58, 59*). Indeed, emotions like the feeling of pleasantness feature prominently in theories of aesthetics (*46, 60-62*), and pleasantness accounts for a substantial share of variance in beauty ratings (*63, 64*). Yet beauty and pleasure are not the same. Our results show that the neural correlates of beauty on the category level are not accounted for by the pleasantness of that category (which we controlled for in our analysis), demonstrating a neural computation of beauty in the visual system that goes beyond the coding of emotional associations with an object category.

Beyond affective factors, there is a wealth of semantic associations for objects of a certain category, which can modulate our aesthetic experiences (*65*) and which might also modulate visual brain responses to these categories. Our results suggest that semantic associations are not critical for the neural correlates of beauty to emerge. First, beauty-related neural responses emerged very rapidly, and in a rapid presentation regime that prevented prolonged cognitive engagement with the stimulus. Second, we show that neural activity recorded from monkey visual cortex predicts beauty, too. This shows that even in the absence of language and the semantic associations transported by it, visual representations are indicative of human beauty judgments.

While our study demonstrates a spontaneous representation of beauty across object categories, it prompts further questions on the nature of the underlying representations. First, while our study suggests that visual feature coding is critical for distinguishing the beauty of categories, it remains unclear which features render a category beautiful. Future studies could distill the contributions of individual image features, such as contour and curvature (*66*), order and complexity (*67*), meaningful object dimensions (*21*), or behavioral factors like affordances (*40*). Alternatively, future work could focus on overarching computational principles that transcend individual feature preferences, such as the efficient allocation of metabolic processing demands (*15*), the efficient integration of visual features (*14*) or the spatiotemporal predictability of stimulus features (*45*). Second, we quantified the beauty of object categories by averaging ratings across observers, and we averaged neural responses across participants. This approach revealed robust neural representations of object beauty, measured as mean rated beauty across individuals, thereby indicating a level of universality. Yet, aesthetic appeal is known to differ between individuals (*68-70*), with traits like beauty appreciation (*71*) or taste typicality (*72*) influencing individual preferences. Future studies could combine neural recordings and subsequent beauty ratings in the same participants to test whether even the earliest neural computations of beauty already reflect individual preferences.

Our results revealed that neural activity in the macaque ventral visual stream may predict human preferences in beauty. This raises the question whether the animals would show similar preferences to humans, for example when probed in a preferential looking or a self-paced free viewing paradigm. Such findings would suggest an underlying universality in the perception of beauty, which may fundamentally stem from commonalities in visual processing across species. Such commonalities may arise from a preference for natural feature distributions, where patterns reverberating an intact nature, such as for example in biophilic design (*73, 74*), may be ones to commonly entice the visual system.

## Materials and Methods

### Visual stimuli: THINGS

In the present study, we aimed to investigate the spatiotemporal neural dynamics of object category beauty. To broadly sample visual object categories, we used images from the THINGS database (*21*). The database consists of 1,854 object categories, each represented by ≥12 image exemplars (in total 26,107 images). These were extended by another 1,854 THINGSplus exemplar images by Stoinski, Perkuhn and Hebart (*26*). All images are high-quality photographs, depicting concrete objects or other nameable categories in a naturalistic environment, from animals and plants over food and clothing items to a variety of manmade objects and nameable materials (e.g., gold, wood, sand). The exemplar images may display one or more individual objects pertaining to their category (e.g., multiple apples or flowers).

### Object beauty ratings

To determine the perceived beauty of each of the 1,854 categories in THINGS, we recruited workers via *Amazon Mechanical Turk* (AMT). Before ratin, informed consent was obtained from all participants. After non-adherent worker exclusion (at least three HITs of either a total duration below 30s on 10 subsequent trials, a variance in rating of less than 1 on 10 subsequent trials, or both), *N*=3,750 (1,895 female, 1,810 male, 45 other or unknown) workers (Age: *M*=38.26, *SD*=11.07, range: 18-80), rated the first six exemplars of all 1,854 object concepts as well as all THINGSplus exemplars, totaling 12,978 images. By sampling multiple images per category, we aimed to eliminate differences arising from particularly beautiful images of any given category, ensuring representative category-level estimates. Workers were asked to rate the beauty of the presented image on a 9-point Likert scale (1 = not beautiful at all, to 9 = very beautiful). For each image, *M*=30.51 ratings (*SD*=1.58, range: 24-36) were acquired. The collected data was highly reliable, showing Spearman-Brown corrected split-half reliability of *r*=.91. In addition, ratings were highly similar within categories: Observing the standard deviation across the 7 images per category, the 1,854 categories showed a mean SD of 0.65 (*SD*=0.22, range: 0.12-1.64), on the 1-9 scale. To obtain category-level beauty estimates, ratings were averaged across all exemplar images within each object category. After averaging, the standard deviation across all 1,854 category-level beauty values was 1.07 (*M*=5.04, range: 2.57-8.10). As an overview, the Top 10 most and least beautiful object categories are displayed in Supplementary Figure 3. For similarity-based analyses, we constructed Representational Dissimilarity Matrices (RDMs) (*75*), where the absolute difference in mean beauty rating between two categories served as the measure of dissimilarity.

### Object property ratings

To reduce the potential confounding effects of other object attributes, we included a set of 11 control ratings taken from THINGSplus (*26*), which cover ratings of manmadeness, naturalness, animacy, heaviness, preciousness, pleasantness, arousal level, as well as an objects’ ability to move, to be moved, to be grasped, and to be held. These ratings were obtained through AMT in a way similar to the beauty ratings (*26*). The mean ratings for all object properties were downloaded via Open Science Framework (OSF, https://osf.io/jum2f/). Furthermore, we also included “cuteness” ratings collected on AMT (*M*=29.45 ratings per image, *SD*=2.70, range: 21-39), which were acquired for the purpose of another study (*76*). In total, 12 control ratings were included as covariates in all analyses (see below). For this purpose, RDMs were constructed based on absolute pairwise differences between ratings, identical to the beauty RDMs.

### Human EEG data

EEG recordings were taken from the THINGS EEG2 dataset (*16*). In the experiment underlying the dataset, *N*=10 healthy adults (mean age 28.5 years, 8 female, 2 male) viewed all 1,854 object concepts. The authors split the concepts into a test partition of 200 categories, with 1 image per category and 80 image repetitions, and a training partition of 1,654 categories, with 10 images per concept and 4 image repetitions. Only the training partition was chosen for the present study. In this partition, participants viewed the first 10 exemplars of 1,654 categories, in a rapid serial visual presentation (RSVP; also see) paradigm (100 ms stimulus, 100 ms fixation), with a visual angle of 7°. Each image was repeated four times. Participants performed a detection task, indicating – after a rapid sequence of 20 images – whether that sequence contained a picture of *Buzz Lightyear*, the “Toy Story” character. The EEG signal was recorded on 63 EEG channels at a sampling rate of 1000 Hz, with online filtering from 0.1 Hz to 100 Hz and referencing to the Fz electrode of the international 10-10 system. The raw EEG data was downloaded via OSF (https://osf.io/3jk45/) alongside the accompanying offline preprocessing script, which was downloaded via GitHub (https://github.com/gifale95/eeg_encoding) and developed by Gifford, Dwivedi, Roig and Cichy (*16*) using the MNE package (*77*) for Python. This preprocessing script was subsequently adapted and used to generate the final dataset for the analysis, using Python 3.12.1 and MNE-Python 1.6.1. During preprocessing, the raw EEG data was epoched from -100ms to +700ms relative to stimulus onset, baseline correction was applied to individual trials and channels, the data was downsampled to 200 Hz, and multivariate noise normalization was applied (*16*).

Before further analysis, the data was exported to Matlab R2023a (The MathWorks Inc., Natick, MA, USA). The four repetitions for each image were averaged, then trials were averaged across exemplars for every category, yielding a data matrix with 1,654 trials (i.e., average trials for each category), 63 channels and 161 timepoints.

Neural RDMs were constructed using the Matlab toolbox CoSMoMVPA (*78*). Firstly, the dimensionality of the data was reduced by means of Principal Component Analysis (PCA) (*79*), keeping the number of components explaining 99% of variance at each timepoint. For our main analysis, the data was then trimmed to the subset of 634 object categories present across all neural data sources (EEG, MEG, fMRI). The data was then converted into RDMs by calculating the Pearson distance (1 minus the Pearson correlation) for each pairwise comparison and subsequently vectorized by converting the lower triangle of each RDM (excluding the central diagonal, where a given datapoint is paired with itself) into a column vector, subsequently called Representational Dissimilarity Vector (RDV). The set of 634 remaining object concepts produced RDVs with a length of 200,661 elements (i.e., unique pairwise comparisons). Lastly, for every timepoint, the EEG RDVs were averaged across subjects.

### Human MEG data

In supplementary analyses, we also replicated our findings with MEG recordings, using the THINGS-data MEG dataset (*17*). Here, *N*=4 healthy adults (mean age 23.25 years, 2 female, 2 male) viewed the first 12 exemplars of all 1,854 object concepts in fast succession (500ms stimulus, 1000±200ms fixation), with a visual angle of 10°. During viewing, participants performed an oddball detection task, in which they were asked to detect a synthetic image not depicting an object. MEG data was acquired with 275 axial first-order gradiometers at 1200 Hz. Data preprocessing and cleaning consisted of bandpass filtering between 0.1 and 40 Hz, epoching from -100ms to 1300ms relative to stimulus onset, baseline correction, and subsequent downsampling to 200 Hz, matching the EEG dataset. Because of one noisy and three dysfunctional channels, data was available for 271 channels. The preprocessed data for each participant was downloaded via figshare (https://doi.org/10.25452/figshare.plus.c.6161151.v1), however participants 1 and 4 were excluded from the dataset due to concerns about data quality caused by insufficient attention during data collection (see (*17*); also see Supplementary Figure 5 for participant-level RSA results). The datasets of participants 2 and 3 were cropped to epochs of -100ms to 700ms, identical to the EEG dataset, then the oddball trials were removed, and lastly, trials were again averaged across image exemplars, yielding a matrix of 1,854 trials (category-level), 271 channels, and 161 timepoints. In a procedure identical to the EEG data (PCA, then Pearson distance), we then computed RDMs, vectorized them into RDVs, and averaged them across subjects.

### Human fMRI data

fMRI recordings were taken from the THINGS fMRI dataset (*17*). In the experiment underlying the dataset, *N*=3 healthy adults (mean age: 25.33 years, 2 female, 1 male) viewed the first 12 exemplars of 720 object categories. Object images were presented in the identical experimental paradigm as the MEG data, albeit at a slower pace (500ms stimulus, 4000ms fixation). Data was acquired using a 3 Tesla scanner and a 32-channel head coil. Whole-brain functional MRI data was collected with 2mm isotropic resolution. Preprocessed and denoised fMRI data, in the form of single-trial BOLD response estimates, were downloaded via figshare (https://doi.org/10.25452/figshare.plus.c.6161151.v1). The downloaded data was imported into Matlab and catch trials were removed, resulting in 8,640 image trials. Lastly, we averaged across image exemplars, leaving 720 voxel-wise response estimates (i.e., one per object category).

### Non-human primate multi-unit activity

Neural recordings from monkey visual cortex were taken from the THINGS Ventral-stream Spiking Dataset (TVSD) (*20*). In the experiment underlying the dataset, *N*=2 adult macaque monkeys (males, 6 and 7 years old) viewed the first 12 exemplars of all 1,854 object categories in THINGS, with each image presented a single time. In addition, 100 images were shown 30 times, as a test set for encoding models; these test trials were not included in the analysis. Experimentation was spread across 7 days (monkey 1) and 4 days (monkey 2). Both monkeys were implanted with electrode arrays spanning their V1 (7 arrays in monkey 1 and 8 in monkey 2), V4 (4 arrays in monkey 1 and 3 in monkey 2), and IT cortex (4 arrays in monkey 1 and 5 in monkey 2). MUA recorded from these arrays was downloaded from https://gin.g-node.org/paolo_papale/TVSD. Our main analysis was performed on normalized MUA, where the activity was averaged for each object category in a 100ms time-window centered on the peak response for each region (V1: 25-125ms, V4: 50-150ms, IT: 75-175ms). From these recordings, we created an RDM for each region and both animals by computing pairwise correlations of MUA response vectors across electrode contacts and subtracting the correlations from 1. In Supplementary analyses, we also analyzed the full MUA time courses, where RDMs were created in steps of 10ms.

### Spatial preselection – fMRI RSA

Before conducting the M/EEG-fMRI fusion analyses, we performed a hypothesis-free spatial preselection in the fMRI voxel space. The rationale for this was that brain regions which do not represent beauty in the first place do not need to be included in the subsequent fusion analysis, saving compute time and reducing false-positive rates.

In our preselection routine, we first parcellated the subject-specific brain-wide fMRI data using the homotopic local-global parcellations by Yan et al. (*24*) in their 100-parcel resolution, which were downloaded from GitHub (https://github.com/ThomasYeoLab/CBIG/tree/master/stable_projects/brain_parcellation/Yan2023_homotopic/parcellations/MNI/kong17). The parcellation, originally in MNI152-space, was transformed into individual subject space for each fMRI subject, using inverse transformation files from Hebart, Contier, Teichmann et al. (*17*). Subsequently, for every subject and parcel separately, we performed a reliability-based voxel preselection: Every voxel was ranked by descending noise ceiling, subsequently selecting the best 100 voxels, while ensuring that the 100th voxel did not fall below a 5% cutoff value. This resulted in a final set of 10,000 voxels per participant, across 100 parcels. After trimming down the fMRI category-level responses to the matching 634 object concepts, we constructed 50 RDMs using the 200 voxels of every homotopic bilateral parcel pair, again calculated via Pearson distance, and again converting them into RDVs. Lastly, RDVs were again averaged across subjects.

To preselect the relevant bilateral parcels for the fusion commonality analysis, these RDVs were used in a simple Representational Similarity Analysis (RSA) (*75*), correlating them with the beauty RDVs (Pearson correlation). After statistical testing, activity in 10 bilateral parcels was significantly (FDR-corrected *p*<.05) correlated with beauty (see Supplementary Table 1). We then performed fusion commonality analysis (described below) on these 10 bilateral parcel pairs, for all EEG (or MEG, see Supplementary Fig. 1) timepoints.

### Similarity-based fusion commonality analysis

To track how spatiotemporal brain responses relate to the perceived beauty of object categories, we performed a model-based M/EEG-fMRI fusion approach (*18, 22, 23*), inspired by (*19*), in which we used commonality analysis (*25*) to isolate the unique contribution of beauty to the shared representations between these modalities. For each time point (EEG/MEG) and location (fMRI parcel), we computed the semipartial correlation between neural RDVs, controlling for the influence of all object properties except beauty, effectively removing the contribution of those properties from the relationship between the spatial and temporal brain responses. We again computed the semi-partial correlation between the neural RDVs, partialling out all object properties, but now also including beauty, the predictor of interest. At each timepoint and location, both correlation coefficients were squared, yielding coefficients of determination (R^2^), and the second coefficient was subtracted from the first. Hence, the difference between the two values is the commonality, the shared variance between EEG/MEG and fMRI explained uniquely by the predictor of interest – the beauty RDV – when partialling out all additional control variables. The resulting commonality score (arbitrary unit) thus reflects the unique contribution of object category beauty to the spatiotemporal neural data in similarity space. We describe this commonality *C* formally at timepoint *t* and location *p* (parcel) as:

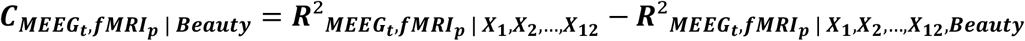

where *X* represents a control variable. We also ran a control analysis in which we exchanged the fMRI RDVs and the M/EEG RDVs as dependent and independent variables. This yielded highly similar results.

### RSA on non-human primate MUA

MUA in monkey V1, V4, and IT was analyzed in an RSA framework. We vectorized the RDM for each region and animal into an RDV and then tested whether these RDVs were predicted by the beauty RDVs. We did this in a partial correlation analysis, where we correlated the MUA RDVs with the beauty RDV while partialling out the 12 control variables (i.e., the other object property ratings, analogous to the EEG-fMRI fusion). This yielded a partial correlation for each region and animal, indicating how well the regional activity is predicted by the beauty of object categories, after accounting for the other ratings.

### Statistical testing

Statistical significance was determined via one-sided permutation tests. For the human fMRI RSA and the non-human primate MUA RSA, the neural RDM (i.e., the critical dependent variable) was shuffled randomly for 2,500 iterations per timepoint. Repeating the analyses with shuffled RDMs yielded a null distribution of permuted (partial) correlation coefficients. This modest but adequate (*80*) number of permutations was chosen to keep the compute time feasible, given the large RDM size. The p-value was determined by the proportion of permuted coefficients exceeding the true coefficient. All gathered p-values of an analysis were simultaneously corrected for multiple comparisons using the False Discovery Rate (FDR) approach, specifically the Benjamini & Hochberg (*28*) method, implemented in Matlab. For the M/EEG-fMRI fusion commonality analyses, we developed and deployed a novel permutation testing approach to reliably correct for the inflations of commonalities that arise from collinearities between predictors (e.g., the EEG and the human rating predictor of interest). In simple permutation testing, these collinearities would be ignored by default, hence any existing collinearities between predictors would typically result in highly significant commonality values. We corrected for this by preserving collinearities throughout permutation testing. In the present case of predicting M/EEG RDVs from fMRI and beauty RDVs, we computed the strict lower bound of commonality by independently fitting the predictors to their respective sign-permuted residuals for 2,500 iterations, obtaining 2x2,500 surrogate estimates. From this, we derived a maximum statistic null distribution by selecting the maximal value between either, for every permutation iteration and timepoint separately. The resulting final 2,500 values were then compared to the observed commonality values, separately per timepoint and parcel, and p-values determined as well as FDR-corrected in identical fashion as outlined above.

## Author contributions

PAS: Conceptualization, Formal analysis, Investigation, Methodology, Software, Writing – original draft, Writing – review & editing

RS: Software, Writing – review & editing

MNH: Funding acquisition, Investigation, Methodology, Software, Supervision, Writing – review & editing

DK: Conceptualization, Formal analysis, Funding acquisition, Methodology, Software, Supervision, Writing – original draft, Writing – review & editing

## Data and code availability

Data and code are freely available in the following repository: https://doi.org/10.5281/zenodo.21192832

## References

1. A. L. Knoll et al., Experiencing Beauty in Everyday Life. Scientific Reports 14, 9463 (2024).

2. R. Weigand, T. Jacobsen, Beauty and the busy mind: Occupied working memory resources impair aesthetic experiences in everyday life. PLoS One 16, e0248529 (2021).

3. A. A. Brielmann, N. H. Buras, N. A. Salingaros, R. P. Taylor, What Happens in Your Brain When You Walk Down the Street? Implications of Architectural Proportions, Biophilia, and Fractal Geometry for Urban Science. Urban Science 6, (2022).

4. A. A. Brielmann, D. G. Pelli, Beauty Requires Thought. Current Biology 27, 1506–1513 (2017).

5. T. Ishizu, S. Zeki, Toward a brain-based theory of beauty. PLoS One 6, e21852 (2011).

6. E. A. Vessel, A. I. Isik, A. M. Belfi, J. L. Stahl, G. G. Starr, The default-mode network represents aesthetic appeal that generalizes across visual domains. Proceedings of the National Academy of Sciences of the United States of America 116, 19155–19164 (2019).

7. S. Righi, G. Gronchi, G. Pierguidi, S. Messina, M. P. Viggiano, Aesthetic shapes our perception of every-day objects: An ERP study. New Ideas in Psychology 47, 103–112 (2017).

8. J. Chen, Y. Cheng, The relationship between aesthetic preferences of people for ceramic tile design and neural responses: An event-related potential study. Frontiers in Human Neuroscience 16, 994195 (2022).

9. D. Kaiser, K. Nyga, Tracking cortical representations of facial attractiveness using time-resolved representational similarity analysis. Scientific Reports 10, 16852 (2020).

10. D. Kaiser, Characterizing Dynamic Neural Representations of Scene Attractiveness. Journal of Cognitive Neuroscience 34, 1988–1997 (2022).

11. C. Conwell, D. Graham, C. Boccagno, E. A. Vessel, The perceptual primacy of feeling: Affectless visual machines explain a majority of variance in human visually evoked affect. Proceedings of the National Academy of Sciences 122, e2306025121 (2025).

12. K. Iigaya, S. Yi, I. A. Wahle, K. Tanwisuth, J. P. O’Doherty, Aesthetic preference for art can be predicted from a mixture of low- and high-level visual features. Nature Human Behavior 5, 743–755 (2021).

13. K. Iigaya et al., Neural mechanisms underlying the hierarchical construction of perceived aesthetic value. Nature Communications 14, 127 (2023).

14. S. Nara, D. Kaiser, Integrative processing in artificial and biological vision predicts the perceived beauty of natural images. Science Advances 10, (2024).

15. Y. Tang, W. A. Cunningham, D. B. Walther, Less is more: Aesthetic liking is inversely related to metabolic expense by the visual system. PNAS Nexus 4, pgaf347 (2025).

16. A. T. Gifford, K. Dwivedi, G. Roig, R. M. Cichy, A large and rich EEG dataset for modeling human visual object recognition. NeuroImage 264, 119754 (2022).

17. M. N. Hebart et al., THINGS-data, a multimodal collection of large-scale datasets for investigating object representations in human brain and behavior. eLife 12, (2023).

18. R. M. Cichy, A. Oliva, A M/EEG-fMRI Fusion Primer: Resolving Human Brain Responses in Space and Time. Neuron 107, 772–781 (2020).

19. M. N. Hebart, B. B. Bankson, A. Harel, C. I. Baker, R. M. Cichy, The representational dynamics of task and object processing in humans. eLife 7, (2018).

20. P. Papale, F. Wang, M. W. Self, P. R. Roelfsema, An extensive dataset of spiking activity to reveal the syntax of the ventral stream. Neuron 113, 539–553.e535 (2025).

21. M. N. Hebart et al., THINGS: A database of 1,854 object concepts and more than 26,000 naturalistic object images. PLoS One 14, e0223792 (2019).

22. R. M. Cichy, D. Pantazis, A. Oliva, Resolving human object recognition in space and time. Nature Neuroscience 17, 455–462 (2014).

23. R. M. Cichy, D. Pantazis, A. Oliva, Similarity-Based Fusion of MEG and fMRI Reveals Spatio-Temporal Dynamics in Human Cortex During Visual Object Recognition. Cerebral Cortex 26, 3563–3579 (2016).

24. X. Yan et al., Homotopic local-global parcellation of the human cerebral cortex from resting-state functional connectivity. NeuroImage 273, 120010 (2023).

25. D. R. Seibold, R. D. McPhee, COMMONALITY ANALYSIS: A METHOD FOR DECOMPOSING EXPLAINED VARIANCE IN MULTIPLE REGRESSION ANALYSES. Human Communication Research 5, 355–365 (1979).

26. L. M. Stoinski, J. Perkuhn, M. N. Hebart, THINGSplus: New norms and metadata for the THINGS database of 1854 object concepts and 26,107 natural object images. Behavior Research Methods 56, 1583–1603 (2024).

27. R. Kong et al., Individual-Specific Areal-Level Parcellations Improve Functional Connectivity Prediction of Behavior. Cerebral Cortex 31, 4477–4500 (2021).

28. Y. Benjamini, Y. Hochberg, Controlling the False Discovery Rate: A Practical and Powerful Approach to Multiple Testing. Journal of the Royal Statistical Society. Series B (Methodological) 57, 289–300 (1995).

29. T. Grootswagers, A. K. Robinson, T. A. Carlson, The representational dynamics of visual objects in rapid serial visual processing streams. NeuroImage 188, 668–679 (2019).

30. D. J. Kravitz, K. S. Saleem, C. I. Baker, L. G. Ungerleider, M. Mishkin, The ventral visual pathway: an expanded neural framework for the processing of object quality. Trends in Cognitive Sciences 17, 26–49 (2013).

31. N. Kriegeskorte et al., Matching categorical object representations in inferior temporal cortex of man and monkey. Neuron 60, 1126–1141 (2008).

32. E. A. Vessel, T. Ishizu, G. Bignardi, in The Routledge International Handbook of Neuroaesthetics, M. Skov, M. Nadal, Eds. (Routledge, London, 2022), pp. 103–133.

33. V. S. Ramachandran, W. Hirstein, The science of art: a neurological theory of aesthetic experience. Journal of Consciousness Studies 6, 15–51 (1999).

34. S. Zeki, Inner vision: An exploration of art and the brain. (Oxford University Press, Oxford, 1999).

35. S. Zeki, J. P. Romaya, D. M. Benincasa, M. F. Atiyah, The experience of mathematical beauty and its neural correlates. Frontiers in Human Neuroscience 8, 68 (2014).

36. C. Redies, Combining universal beauty and cultural context in a unifying model of visual aesthetic experience. Frontiers in Human Neuroscience 9, 218 (2015).

37. R. Reber, N. Schwarz, P. Winkielmann, Processing Fluency and Aesthetic Pleasure: Is Beauty in the Perceiver’s Processing Experience? Personality and Social Psychology Review 8, 364–382 (2004).

38. H. Leder, B. Belke, A. Oeberst, D. Augustin, A model of aesthetic appreciation and aesthetic judgments. British Journal of Psychology 95, 489–508 (2004).

39. H. Leder, M. Nadal, Ten years of a model of aesthetic appreciation and aesthetic judgments: The aesthetic episode - Developments and challenges in empirical aesthetics. British Journal of Psychology 105, 443–464 (2014).

40. S. Righi, V. Orlando, T. Marzi, Attractiveness and affordance shape tools neural coding: insight from ERPs. International Journal of Psychophysiology 91, 240–253 (2014).

41. F. Guo, X.-S. Wang, W.-L. Liu, Y. Ding, Affective preference measurement of product appearance based on event-related potentials. Cognition, Technology & Work 20, 299–308 (2018).

42. S. Wang, C. Xu, L. Xiao, A. S. Ding, The Implicit Aesthetic Preference for Mobile Marketing Interface Layout-An ERP Study. Frontiers in Human Neuroscience 15, 728895 (2021).

43. L. Höfel, T. Jacobsen, Electrophysiological indices of processing aesthetics: Spontaneous or intentional processes? International Journal of Psychophysiology 65, 20–31 (2007).

44. K. Iigaya, J. P. O’Doherty, G. G. Starr, Progress and Promise in Neuroaesthetics. Neuron 108, 594–596 (2020).

45. M. Nadal, M. Skov, The sensory valuation account of aesthetic experience. Nature Reviews Psychology 4, 49–63 (2024).

46. A. A. Brielmann, P. Dayan, A computational model of aesthetic value. Psychological Review 129, 1319–1337 (2022).

47. L. Höfel, T. Jacobsen, Electrophysiological indices of processing symmetry and aesthetics: A result of judgment categorization or judgment report? Journal of Psychophysiology 21, 9–21 (2007).

48. S. Nara, L. Becker, L. Hillebrand, R. Xiang, D. Kaiser, Dynamic neural representations of scene beauty are relatively unaffected by stimulus timing and task. Scientific Reports 16, 15217 (2026).

49. C. J. Cela-Conde et al., Sex-related similarities and differences in the neural correlates of beauty. Proceedings of the National Academy of Sciences of the United States of America 106, 3847–3852 (2009).

50. C. J. Cela-Conde et al., Dynamics of brain networks in the aesthetic appreciation. Proceedings of the National Academy of Sciences of the United States of America 110, 10454–10461 (2013).

51. O. Vartanian, V. Goel, Neuroanatomical correlates of aesthetic preference for paintings. NeuroReport 15, 893–897 (2004).

52. B. Calvo-Merino, C. Urgesi, G. Orgs, S. M. Aglioti, P. Haggard, Extrastriate body area underlies aesthetic evaluation of body stimuli. Experimental Brain Research 204, 447–456 (2010).

53. M. Boccia et al., Where does brain neural activation in aesthetic responses to visual art occur? Meta-analytic evidence from neuroimaging studies. Neuroscience and Biobehavioral Reviews 60, 65–71 (2016).

54. A. M. Belfi et al., Dynamics of aesthetic experience are reflected in the default-mode network. NeuroImage 188, 584–597 (2019).

55. E. A. Vessel, G. G. Starr, N. Rubin, Art reaches within: aesthetic experience, the self and the default mode network. Frontiers in Neuroscience 7, 258 (2013).

56. S. Kühn, J. Gallinat, The neural correlates of subjective pleasantness. NeuroImage 61, 289–294 (2012).

57. T. K. Pegors, J. W. Kable, A. Chatterjee, R. A. Epstein, Common and unique representations in pFC for face and place attractiveness. Journal of Cognitive Neuroscience 27, 959–973 (2015).

58. C. Gao, S. Ajith, M. V. Peelen, Object representations drive emotion schemas across a large and diverse set of daily-life scenes. Communications Biology 8, 697 (2025).

59. R. D. Lane et al., Neuroanatomical correlates of pleasant and unpleasant emotion. Neuropsychologia 35, (1997).

60. M. Skov, M. Nadal, The Nature of Beauty: behavior, cognition, and neurobiology. Annals of the New York Academy of Sciences 1488, 44–55 (2021).

61. I. Biederman, E. A. Vessel, Perceptual Pleasure and the Brain. American Scientist 94, 249–255 (2006).

62. A. K. M. Rezaoul Karim, M. J. Proulx, A. A. de Sousa, L. T. Likova, Do we enjoy what we sense and perceive? A dissociation between aesthetic appreciation and basic perception of environmental objects or events. Cognitive, Affective, & Behavioral Neuroscience 22, 904–951 (2022).

63. O. Vartanian et al., Impact of contour on aesthetic judgments and approach-avoidance decisions in architecture. Proceedings of the National Academy of Sciences of the United States of America 110, 10446–10453 (2013).

64. A. A. Brielmann, A. Nuzzo, D. G. Pelli, Beauty, the feeling. Acta Psychologica 219, 103365 (2021).

65. U. Kirk, M. Skov, O. Hulme, M. S. Christensen, S. Zeki, Modulation of aesthetic value by semantic context: an fMRI study. NeuroImage 44, 1125–1132 (2009).

66. E. G. Chuquichambi et al., How universal is preference for visual curvature? A systematic review and meta-analysis. Annals of the New York Academy of Sciences 1518, 151–165 (2022).

67. E. Van Geert, J. Wagemans, Order, complexity, and aesthetic appreciation. Psychology of Aesthetics, Creativity, and the Arts 14, 135–154 (2020).

68. E. A. Vessel, N. Rubin, Beauty and the beholder: highly individual taste for abstract, but not real-world images. Journal of Vision 10, 18 11-14 (2010).

69. T. Jacobsen, Beauty and the brain: culture, history and individual differences in aesthetic appreciation. Journal of Anatomy 216, 184–191 (2010).

70. H. Leder, J. Goller, T. Rigotti, M. Forster, Private and Shared Taste in Art and Face Appreciation. Frontiers in Human Neuroscience 10, 155 (2016).

71. R. Diessner, Understanding the Beauty Appreciation Trait. (Palgrave Macmillan, Cham, 2019).

72. Y. C. Chen et al., “Taste typicality” is a foundational and multi-modal dimension of ordinary aesthetic experience. Current Biology 32, 1837–1842 e1833 (2022).

73. W. Zhong, T. Schröder, J. Bekkering, Biophilic design in architecture and its contributions to health, well-being, and sustainability: A critical review. Frontiers of Architectural Research 11, 114–141 (2022).

74. K. Gillis, B. Gatersleben, A Review of Psychological Literature on the Health and Wellbeing Benefits of Biophilic Design. Buildings 5, 948–963 (2015).

75. N. Kriegeskorte, M. Mur, P. Bandettini, Representational similarity analysis - connecting the branches of systems neuroscience. Frontiers in Systems Neuroscience 2, 4 (2008).

76. L. M. Stoinski, T. Konkle, M. N. Hebart, Principles of coarse-scale functional organization in occipitotemporal cortex. bioRxiv, (2026).

77. A. Gramfort et al., MEG and EEG data analysis with MNE-Python. Frontiers in Neuroscience 7, 267 (2013).

78. N. N. Oosterhof, A. C. Connolly, J. V. Haxby, CoSMoMVPA: Multi-Modal Multivariate Pattern Analysis of Neuroimaging Data in Matlab/GNU Octave. Frontiers in Neuroinformatics 10, 27 (2016).

79. T. Grootswagers, S. G. Wardle, T. A. Carlson, Decoding Dynamic Brain Patterns from Evoked Responses: A Tutorial on Multivariate Pattern Analysis Applied to Time Series Neuroimaging Data. Journal of Cognitive Neuroscience 29, 677–697 (2017).

80. A. M. Winkler, G. R. Ridgway, G. Douaud, T. E. Nichols, S. M. Smith, Faster permutation inference in brain imaging. NeuroImage 141, 502–516 (2016).

