## Supplement for "Primate visual cortex spontaneously computes the beauty of objects"

| **Parcel No.** (LH, RH) | **Network** [LH (, RH)]* *(24)* | **Area** *(24)* | **MNI152, LH** | | | **MNI152, RH** | | | **AAL annotation** [LH(, RH)]* |
|---|---|---|---|---|---|---|---|---|---|
| | | | **R** | **A** | **S** | **R** | **A** | **S** | |
| 1, 51 | DefaultC | PHC | -28 | -30 | -22 | 28 | -26 | -22 | Fusiform *(next to PHC)* |
| 2, 52 | DefaultC | pCun | -8 | -70 | 46 | 10 | -68 | 46 | Precuneus |
| 10, 60 | DefaultA, ContB | IPL | -42 | -72 | 38 | 48 | -64 | 38 | Angular |
| 32, 82 | DorsAttnB | SPL | -16 | -54 | 66 | 16 | -52 | 68 | Precuneus, Parietal Sup |
| 33, 83 | DorsAttnA, DorsAttnB | PostC | -60 | -28 | 34 | 60 | -22 | 38 | SupraMarginal *(next to PostC)* |
| 34, 84 | DorsAttnA | SPL | -24 | -70 | 46 | 28 | -68 | 48 | Parietal Sup, Occipital Sup |
| 47, 97 | VisualB | 4 | -24 | -68 | -12 | 24 | -66 | -10 | Fusiform |
| 48, 98 | VisualA | 1 | -30 | -88 | 16 | 36 | -84 | 18 | Occipital Mid |
| 49, 99 | VisualA | 2 | -44 | -74 | -10 | 46 | -68 | -12 | Occipital Inf |
| 50, 100 | VisualA | TempOcc | -48 | -68 | 10 | 52 | -62 | 10 | Temporal Mid |

***Supplementary Table 1.*** *Selected bilateral homotopic parcel pairs after RSA with Yan atlas (24) (100-parcel resolution) numbers & annotations, bilateral centroid coordinates in MNI space, and AAL annotations based on given MNI coordinates; Ordered by parcel number. Abbreviations: LH = Left Hemisphere, RH = Right Hemisphere; Default = Default, Cont = Control, DorsAttn = Dorsal Attention; PHC = Parahippocampal Cortex, pCun = Precuneus, IPL = Inferior Parietal Lobule, SPL = Superior Parietal Lobule, PostC = Postcentral, TempOcc = Temporal Occipital; Inf: Inferior, Mid: Middle, Sup: Superior. *[LH(, RH)]: If annotations of LH and RH match, only one is written, otherwise LH is followed by RH.*

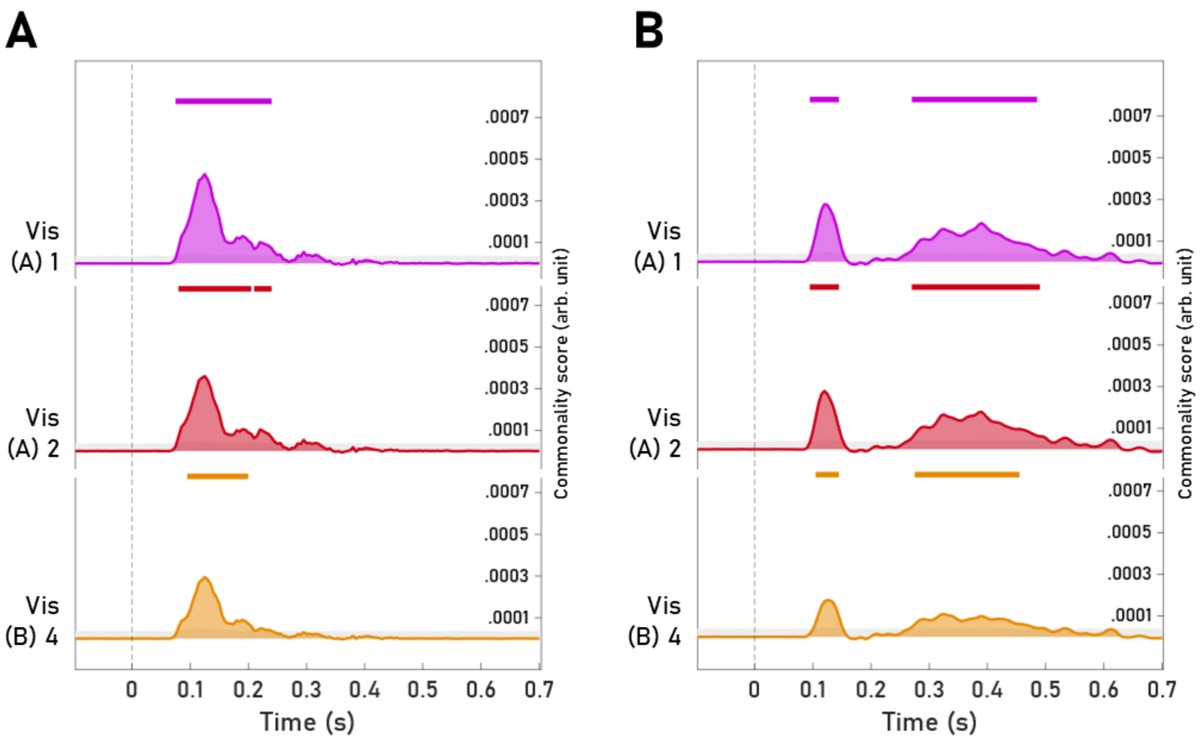


***Supplementary Figure 1. Fusion commonality analyses validations on identical scale to Fig. 1 C and consistent color-coding for direct comparison. A) Validation analysis across exemplar images.*** *This fusion spanned the same 634 object concepts (Fig. 1) but was reduced to images that were not rated for their beauty in the online experiment (trial reduction of both EEG and fMRI data by ~60%), as well as to the three parcels of interest. Significant timepoints (FDR-corrected p<.05) marked by rectangles, in the corresponding parcel color. Scale axes are identical to the full EEG-fMRI fusion reported in Fig. 1 C for direct comparison. The results of this analysis replicate the previous findings of a strong, early peak of a neural reflection of the beauty of object categories in visual cortical areas. The reduced image set resulted in an expectedly weaker representation of beauty overall, as well as a shortened duration of continuously significant timepoints, while the time course remained qualitatively similar.* ***B) Validation analysis across neuroimaging modalities.*** *Replication of main results in Fig.1 C using MEG-fMRI fusion, keeping the underlying stimulus set and preprocessing aligned with the EEG data (see Materials and Methods). Overall, commonality coefficients are broadly lower than in the EEG-fMRI results, likely resulting from the reduced number of participants and image trials. Significant timepoints emerge slightly later (~90ms, up from ~65ms), possibly due to the higher Stimulus Onset Asynchrony in the MEG data. Similarly to the EEG-fMRI results, a uniform pattern with a strong early peak (~120ms) is observed across the three parcels in the visual cortex, while other parcels mainly show weaker early peaks and often stronger, sustained responses after 250ms. The relatively high commonality score in visual parcels between 250ms and 500ms may be a result of subject idiosyncrasy (compare participant-level RSA on MEG data in Supplementary Fig. 5).*

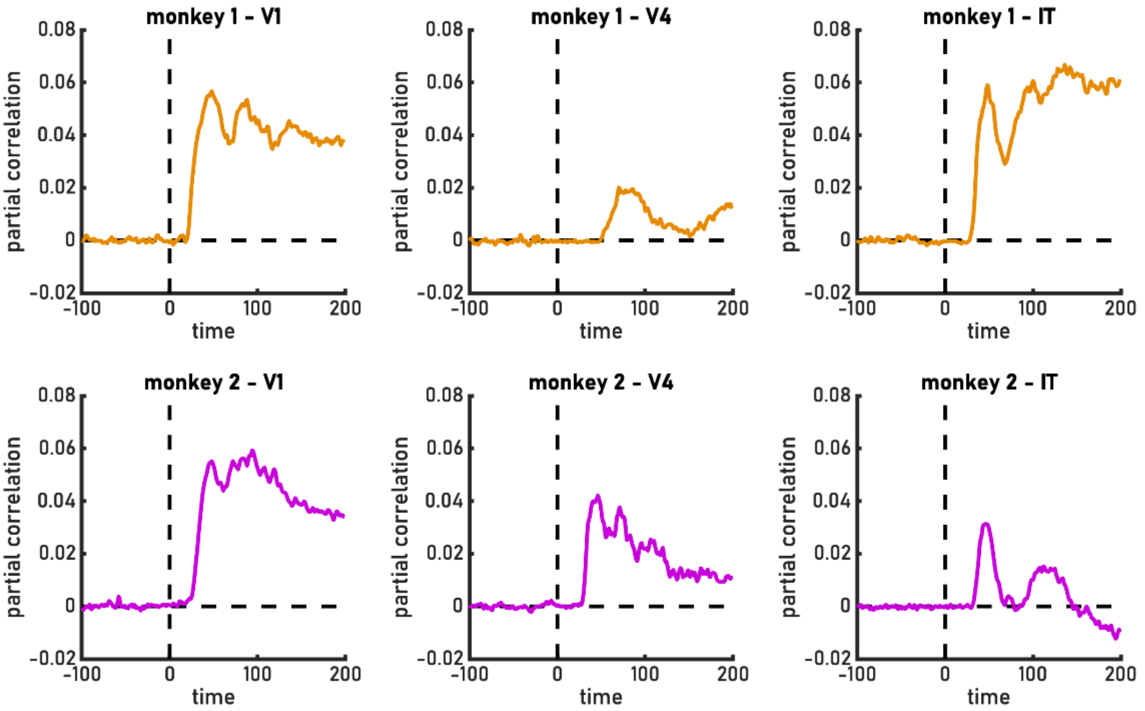


***Supplementary Figure 2.*** *Time-resolved representational similarity analysis on MUA data from monkey V1, V4, and IT. Across both monkeys and all regions, neural activations emerging within 100ms of processing were correlated with perceived beauty. Notably, qualitative differences in the time-resolved results increase with both time (after an early first peak) and in the latter stages of the visual stream hierarchy (V4, IT). This is mirrored by the human participant-level EEG and MEG results in Supplementary Figures 4/5, where idiosyncrasy increases substantially from around 200ms onwards.*

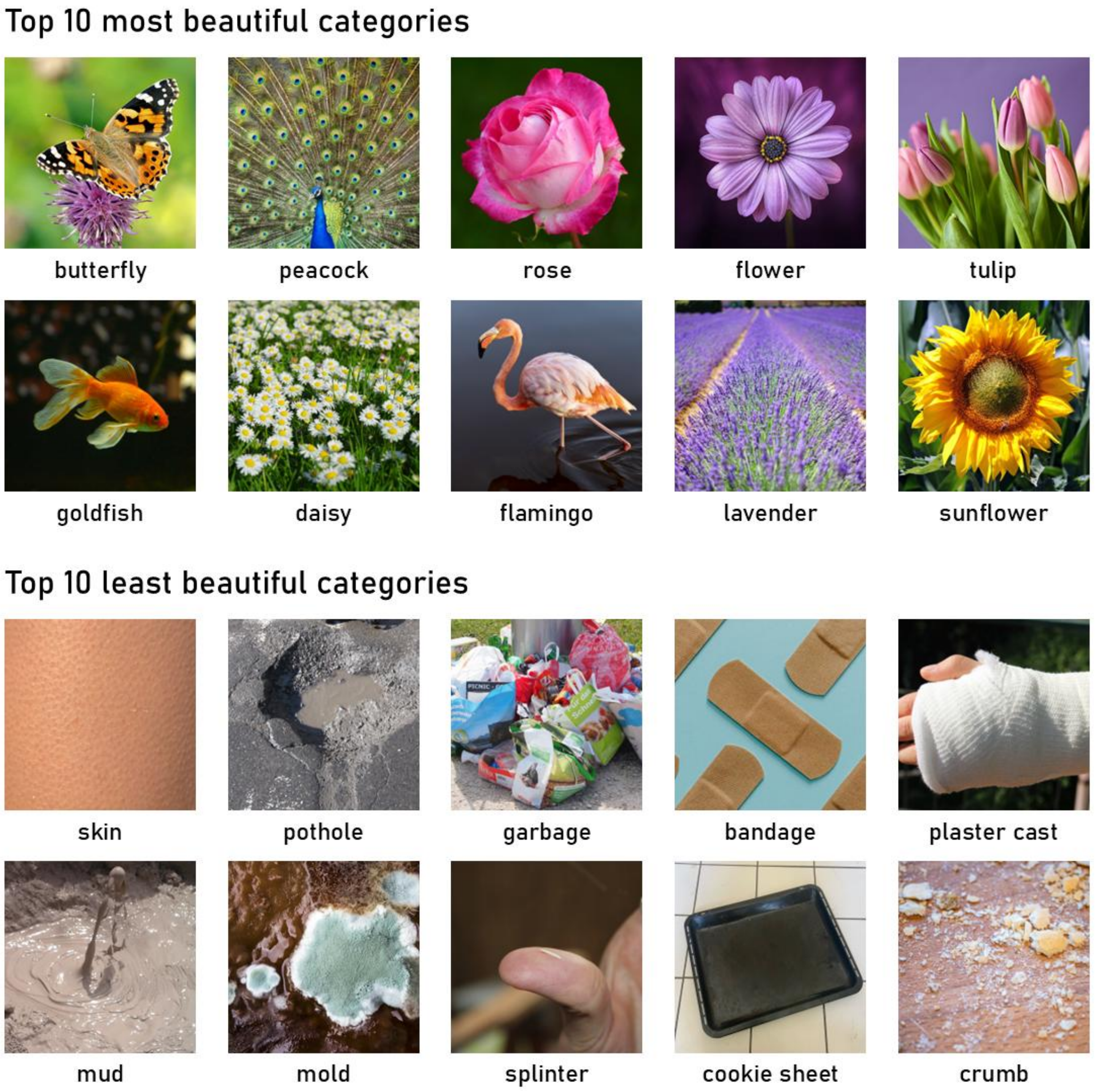


***Supplementary Figure 3.*** *Top 10 most and least beautiful categories across the entire set of 1,854 object categories, by mean beauty across all raters and all rated images. Image exemplars displayed are the license-free THINGSplus (26) images.*

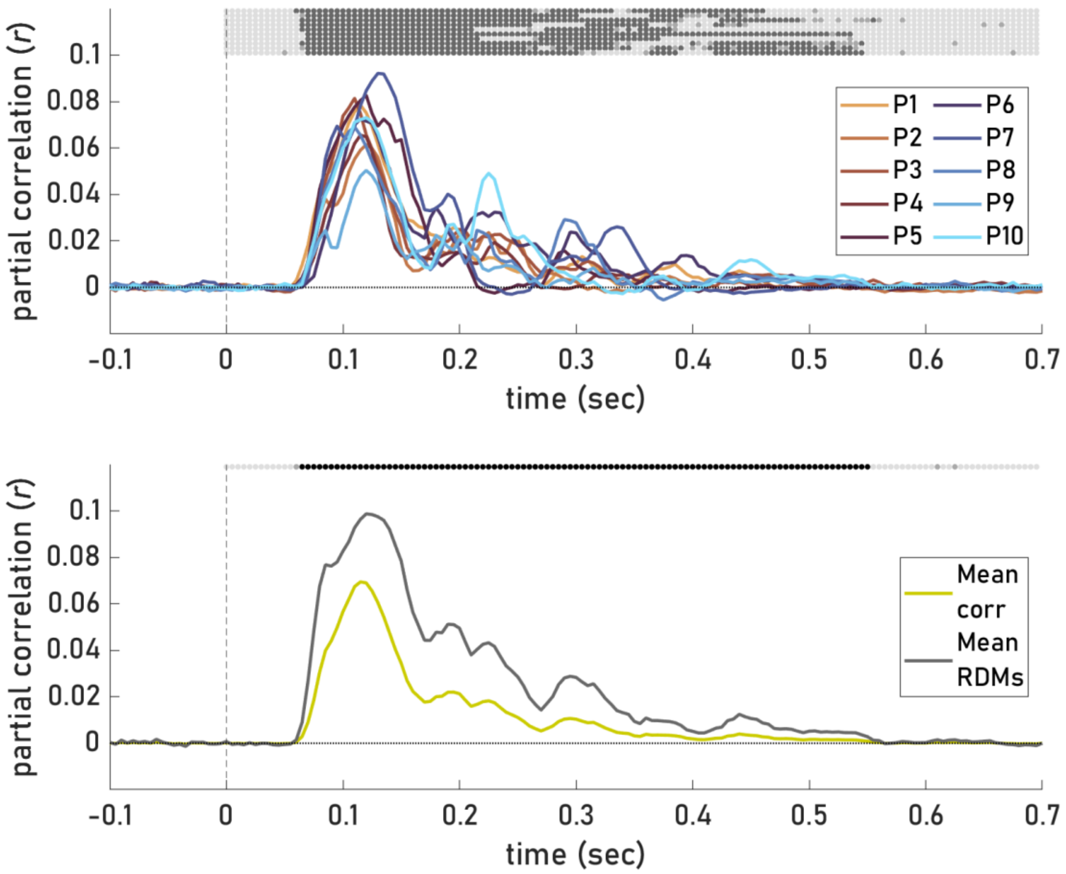


***Supplementary Figure 4.*** *Time-resolved RSA results on beauty correlated with EEG data, when partialling out all predictors, on 1,654 object categories. Individual results above; Below results using mean neural RDMs (similar to main EEG-fMRI fusion; there pruned to 634 object categories), compared to mean partial correlation of individual RSA results. Significance of timepoints displayed via dots (FDR-corrected p<.05 medium gray, p<.01 dark grey) for all 10 subjects individually (top to bottom = 1 to 10).*

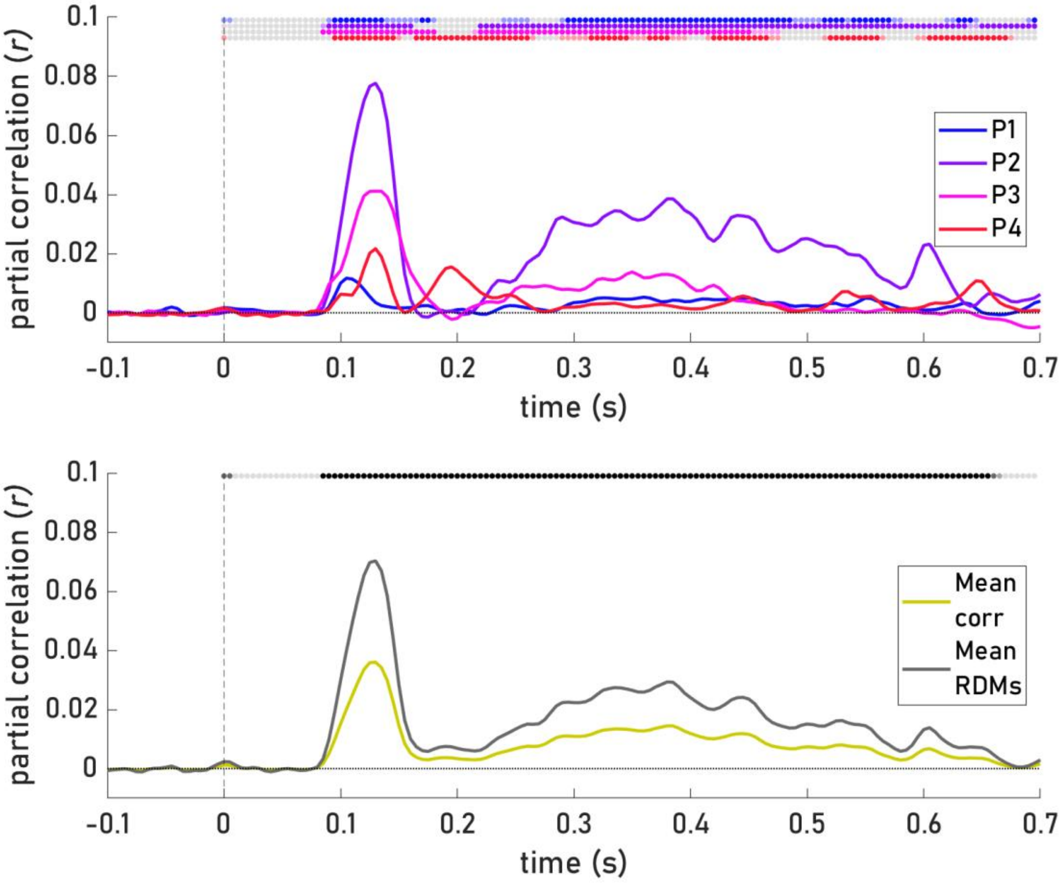


***Supplementary Figure 5.*** *Time-resolved RSA results on beauty correlated with MEG data, when partialling out all predictors, on 1,854 object categories. Individual results above; Below results using mean neural RDMs (mean neural RDMs of P2 and P3 were used in MEG-fMRI fusion, pruned to 634 object categories), compared to mean partial correlation of individual RSA results. Significance of timepoints (FDR-corrected p<.05 in light shade, p<.01 in darker shade) displayed via dots for all 4 subjects individually (in corresponding colors).*
